# Associating Transcription Factors to Single-Cell Trajectories with DREAMIT

**DOI:** 10.1101/2023.06.08.544214

**Authors:** Nathan D Maulding, Lucas Seninge, Joshua M. Stuart

## Abstract

Trajectory methods have enabled the organization of cells into contiguous cellular changes from their transcriptional profiles measured by single cell RNA sequencing. Few methods enable investigating the implied gene regulatory network dynamics from the cell transitions between and along trajectory branches. In particular, there remains an opportunity to develop methods that leverage the predicted “pseudotime” orderings of cells to reveal transcription factor (TF) dynamics. Here we present DREAMIT (**D**ynamic **R**egulation of **E**xpression **A**cross **M**odules in **I**nferred **T**rajectories), a novel framework developed to detect patterns of TF activity along single-cell trajectory branches. It detects significant TF-target associations using a relational enrichment approach. Using a benchmark representing several different tissues, the method was found to have increased tissue-specific sensitivity and specificity over competing approaches. To illustrate the utility of the approach, we apply it to the analysis of a peripheral blood mononucleocyte dataset and discuss several examples of TF networks associated with monocytes and erythrocytes that reveal potential causal relationships among TFs. In summary, DREAMIT provides a useful tool for uncovering potential TF-to-target gene regulatory mechanisms associated with the cell-to-cell transitions predicted by trajectory inference methods.

## Background

A cell’s type and state are a product of gene regulatory mechanisms controlled by transcription factors. Recent advances in single-cell sequencing and transcriptomic methods have allowed researchers to examine gene regulatory networks with high sensitivity and specificity, and “cell trajectory” inference methods have allowed for the inference and identification of transitions between cell states [1,2]. Trajectory methods identify changes in development, maturation, or response to environmental queues using gene expression changes across cells having similar transcriptomes. The dynamic regulation of genes can then be inferred by following their relative expression across cells along a trajectory “branch” transitioning from one cell state to another (Figure 1A).

**Figure 1.**
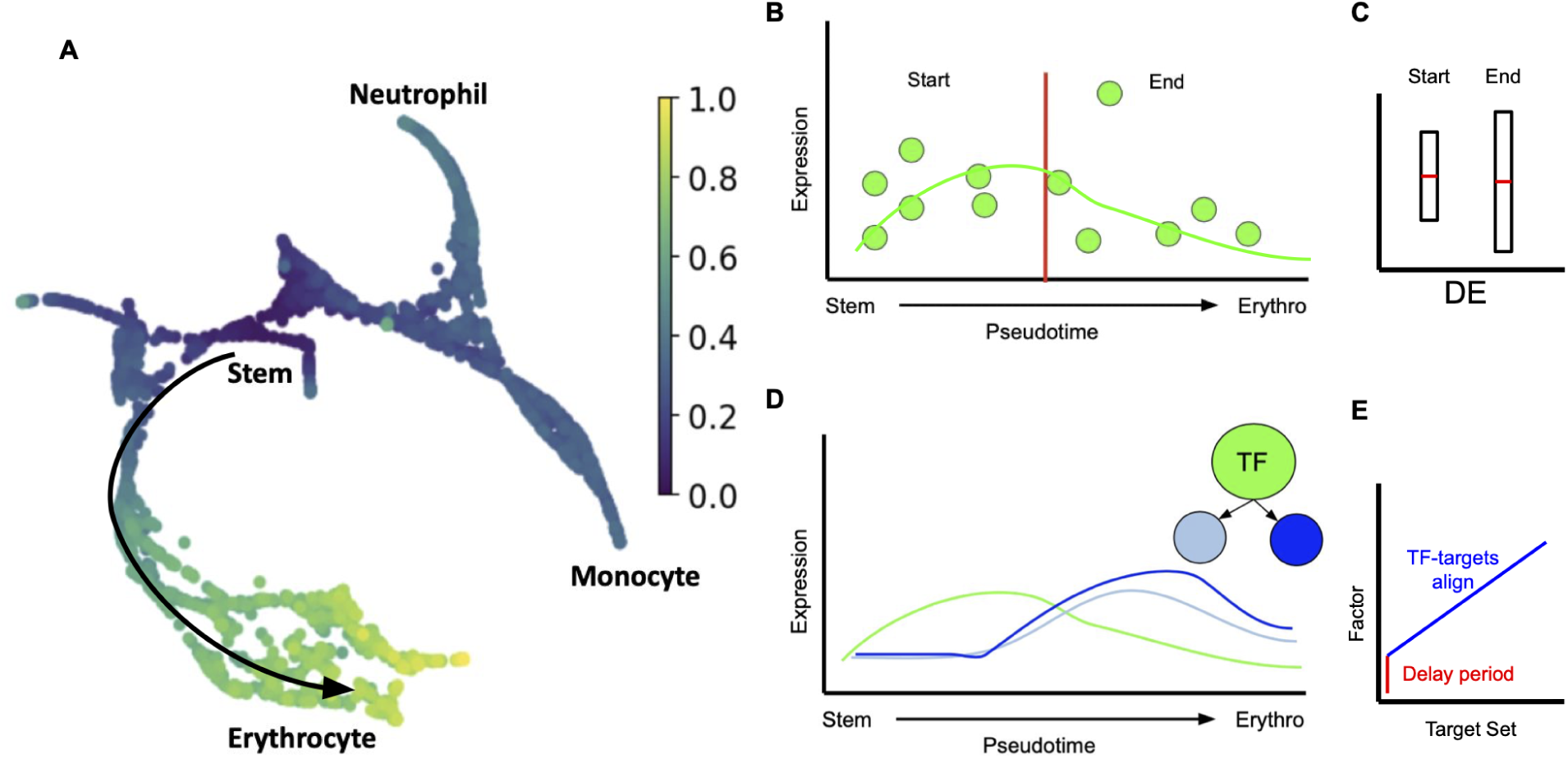
Associating transcription factors (TFs) to trajectory branches via identification of TF-to-target coexpression along pseudotime. **A.** Expression across the cell transition in trajectory branches is used by DREAMIT to infer a dynamic view of TF-to-target gene regulation. **B.** Expression levels of a hypothetical gene in individual cells (y-axis) illustrating the division into arbitrary “start” and “end” states along the pseudotime of a theoretical differentiation process from stem cells to differentiated erythrocytes (x-axis). **C.** Differences in the expression level between “start” and “end” states may not exist which may cause Differential Expression (“DE”) approaches to miss other patterns in the data (e.g. concordant fluctuations in the middle of pseudotime). **D.** DREAMIT models the entirety of the expression on the branch and assesses TF-to-target relationships that look for a consistent relationship between the expression levels (y-axis) of a TF (green line) and its target genes (blue lines) along pseudotime (x-axis). **E.** Alignment plot showing one TF (y-axis) aligned to a “typical” target from a target set (x-axis) illustrating how allowing for a lag or delay (red line) can help a metric pick up on an association between a TF and its targets over a subset of pseudotime (blue line) in which all the targets have the same lag in expression relative to the TF.

Discovering dynamic transcription factor regulation of cell states in scRNAseq data, based on trajectory inference results, has been challenging due to multiple factors, including gene expression dropout, stochastic variation, delayed target response, and variability in chromatin accessibility resulting in discrepancies of known factor-to-target regulation [3–5]. Few methods have been developed that leverage the results of trajectory analysis to identify gene regulatory networks responsible for cell fate decisions. Many methods were developed for application to bulk RNAseq, like GENIE3 [6], and few have been tailored for scRNAseq with the ability to leverage “pseudotime” – a predicted temporal ordering of the cells along the branch of an inferred trajectory. To date, the tools that are available for scRNAseq-based analysis include those based on differential expression analysis, clustering, cell-type annotation, and dimensionality reduction [7–9]. Methods like TRADE-Seq [10] and PseudotimeDE [11] allow users to investigate differential gene expression across a trajectory transition (Figure 1B-C). While methods like SINGE [12] use granger causality ensembles to infer candidate regulatory interactions. However, the performance of methods to infer gene regulatory relationships from trajectories remains challenging and is often difficult to evaluate with any objective criterion.

In this work, we present a method called DREAMIT – Dynamic Regulation of Expression Across Modules in Inferred Trajectories – that analyzes dynamic regulatory patterns along trajectory branches to implicate transcription factors (TFs) involved in cell state transitions found in scRNAseq datasets (Figure 1D-E). DREAMIT performs two “focus” steps to increase the signal from noisy data: 1) a pseudotime focusing to identify a reliable subset of contiguous points along a trajectory and 2) a “target focusing” to identify a subset of targets with the strongest association to a particular TF. First, DREAMIT performs *pseudotime focusing*, together with quantizing and spline smoothing, to ensure the data representation used for detecting relationships are sufficiently robust. It then computes pairwise association scores between all targets and a specific TF (e.g. Pearson correlation) from which it applies a *target focusing* step to identify the subset of targets with the highest scores. Finally, DREAMIT uses a Relational Set Enrichment Analysis (RSEA) test to determine the significance of the set of TF-to-target associations compared to a background model made up of arbitrarily selected targets.

We evaluated DREAMIT’s performance in recovering TFs with known relevance to a particular dataset. The evaluation represents a “bronze standard” due to the lack of suitable reference datasets in which the true underlying regulation driving the major differences among the cells is known as has been previously noted [12]. We used a TF-Marker database to assess the ability of DREAMIT to identify TFs with known roles in specific tissues that have previously been reported as high-confidence markers of those tissues [13]. We found that DREAMIT outperformed traditional approaches of differential expression and GENIE3 in a number of cases.

## Results

### DREAMIT Identifies Distinct PBMC markers

To evaluate DREAMIT’s specificity in a highly curated setting in which confident tissue-specific transcription factor (TF) regulation is well known, we chose a peripheral blood mononucleocyte (PBMC) dataset from Paul et. al. [14]. This dataset contains stem cells transitioning to various blood cell types (erythrocytes, monocytes, neutrophil) suitable for trajectory branch inference and contains a well-characterized set of marker genes. We estimated the accuracy of the methods using an average of 16.3 markers per branch by calculating precision-recall, and early precision (see Methods). DREAMIT was found to have the highest average precision and early precision (AUC=0.57, E= 1.00) while the other approaches had lower estimates – smoothGENIE (AUC=0.46, E=0.38), rawGENIE (AUC=0.43, E=0.27), rawDE (AUC=0.33, E=0.29), smoothDE (AUC=0.33, E=0.17), rawDEtargets (AUC=0.49, E=0.40), and smoothDEtargets (AUC=0.32, E=0.17) (Figure 2A).

**Figure 2.**
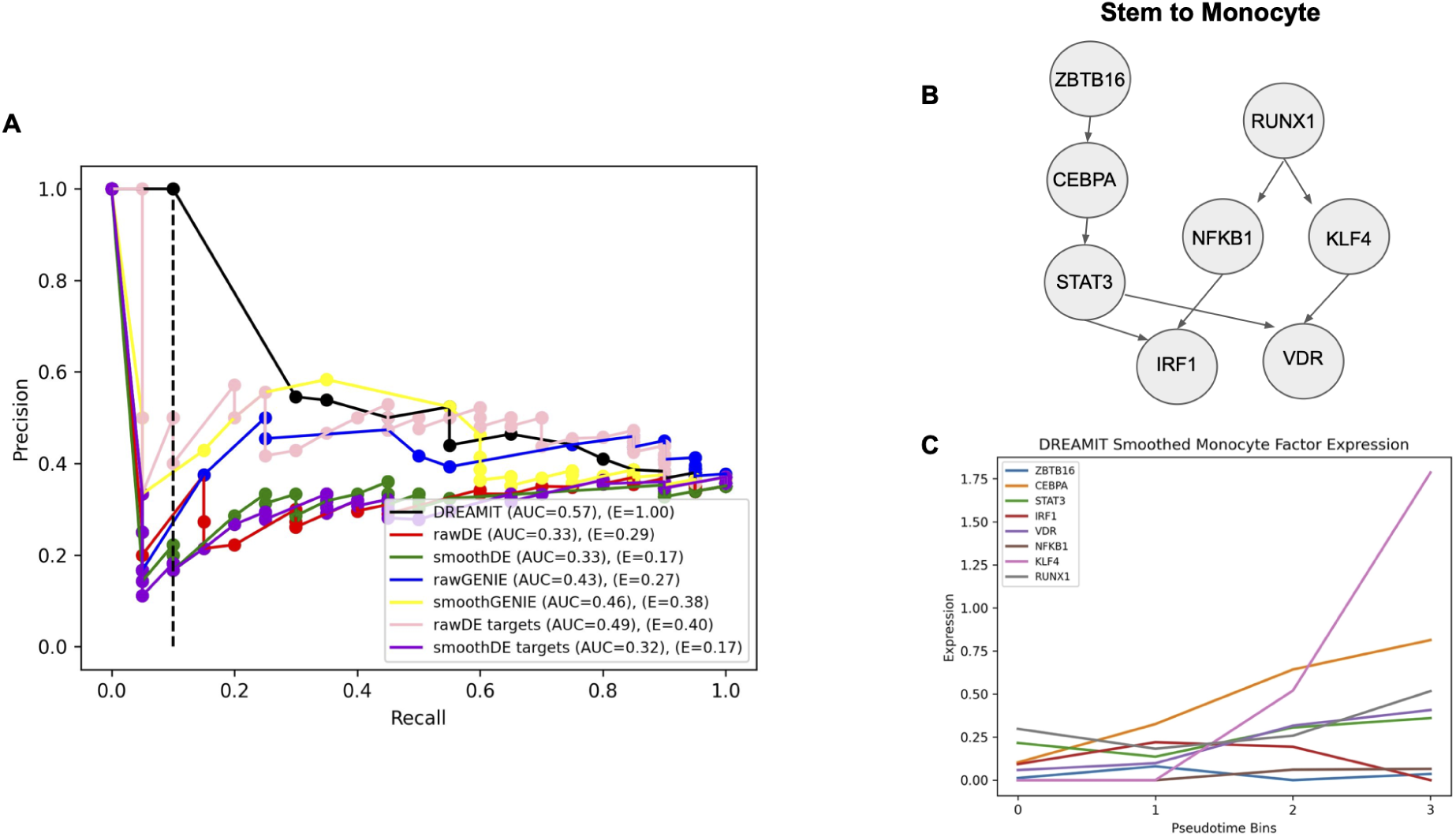
Performance of inferring blood differentiation TFs on the PBMC dataset. **A.** Precision-recall curve measuring rate of identifying blood-related transcription factors annotated in TF-Marker DB from the analysis of the PBMC dataset. A dotted line is plotted on the precision-recall to denote where early-precision is considered. **B.** TF-TF relationships found by DREAMIT depicted for the stem to monocyte branch from this PBMC dataset. **C.** The expression (y-axis) of the TFs from part B plotted across pseudotime (x-axis).

### DREAMIT inference of gene regulatory logic for PBMC fate specification

The majority of TFs found by DREAMIT were annotated as markers of stem or blood, which is consistent given the method had as input trajectories from the Paul et al dataset representing a transition from stem to differentiated blood cell types. For example, on the stem-to-erythrocyte trajectory branch, there were 16 total TFs found to be significant with DREAMIT, five of which were known blood and peripheral blood markers from the TF-marker database (RUNX1, GATA1, EGR1, STAT1, ETS1). Additionally, there were three other factors found that were annotated as stem cell markers (YBX1, MYC, RELA). Of the other 8 TFs reported to be significant by DREAMIT, 6 had literature support linking them to roles in stem and PBMC development – DNMT1 [15], EZH2 [16,17], E2F4 [18,19], KLF6 [20,21], NFE2L2 [22], TP53 [23,24] – while the remaining two TFs, MYB and MYCN, had a less clear relationship. As another example, on the the stem-to-monocyte branch, a total of 13 TFs were reported as significant by DREAMIT, with 6 of them (CEBPA, ETS1, IRF1, ATF4, RUNX1, STAT3) having annotations of blood and peripheral blood markers, and 3 annotated as stem cell markers (IRF8, KLF4, NFKB1). Of the four remaining TFs found on the branch – ZBTB16, MYCN, ELF1, VDR – the vitamin D receptor (VDR) was the only factor with established literature showing its involvement in monocytes [25,26].

To further investigate how DREAMIT findings provide insight to the temporal cascade of effects observed on a trajectory a TF-to-TF network was created. For the significant DREAMIT findings from the high fidelity paul et. al. PBMC dataset [14], TF-to-TF relationships were plotted for the stem-to-monocyte trajectory (Figure 2B). For all TFs found to have a significant association with the branch by DREAMIT, a network was created linking the TF to other TFs if the second TF was also recorded as a target in the TRRUST database. Inspecting the (smoothed) expression of the TFs in the network reveals a concordance with the temporal changes in expression and the connectivity in the TF-to-TF network (Figure 2C). For example, the factors STAT3 and RUNX1 are upregulated at the initial time point and their targets are upregulated in a later time point, consistent with the known regulatory relationships. In another example, a concurrent increase in ZBTB16 and CEBPA (regulated by ZBTB16) was observed followed by increases in their downstream target TFs – STAT3 (regulated by CEBPA) and VDR (regulated by STAT3). On the stem-to-erythrocyte trajectory branch, DREAMIT identified an analogous TF-to-TF network (data not shown). In that case, RUNX1, EGR1, MYCN, and ETS1 were found to increase in the early stages of the trajectory branch with expression increases across the TF network in later stages.

### DREAMIT Identifies tissue-relevant TFs at a higher rate than standard approaches

To systematically estimate the accuracy of a method’s ability to implicate TFs for a given trajectory branch, one requires a variety of tissues with many trajectory branches where the role of TFs have been well documented. However, single cell analyses for most tissues have not been performed to determine a set of reliable TFs. Even so, some information has been collected in repositories like TF-Marker DB [13]. To this end, we collected 7 datasets for which at least one TF was recorded in both TRRUST and TF-Marker DB and was annotated as regulating cells in tissues assayed by the experiments. In total, this “bronze standard” benchmark contained 84 TFs annotated as relevant (207 instances of a TF associated with a dataset), 15 trajectory branches encompassing 6 different tissues (brain, heart, embryo, retina, bone marrow, testis).

To evaluate the methods’ ability to capture tissue-specific TFs in the benchmark datasets, we used a precision-recall analysis. Precision recall is appropriate in cases like this where we expect many more negatives than positives due to the fact that we treat all of the unknown markers in the benchmark as negatives. For a robust comparison, we considered the aggregate performance of DREAMIT, Differential Expression (DE), and GENIE3 on all branches and tissues. To compare the methods’ performance, we used the area under the curve (AUC) to assess general performance across all recall levels, as well as “early precision” (E) [12] to provide a measure of performance at a stringent level of confidence as only the top-ranked relationships are considered in its calculation.

DREAMIT had the highest average precision (AUC=0.20) surpassing its competitors, rawDE (AUC=0.13), smoothDE (AUC=0.13), rawDEtargets (AUC=0.16), smoothDEtargets (AUC=0.16), rawGENIE (AUC=0.16), and smoothGENIE (AUC=0.18) (Figure 3A). Additionally, DREAMIT demonstrates a much stronger early precision (E=0.42) compared to rawDE (E=0.09), smoothDE (E=0.08), rawDEtargets (E=0.19), smoothDEtargets (E=0.13), rawGENIE (E=0.33), and smoothGENIE (E=0.21).

**Figure 3.**
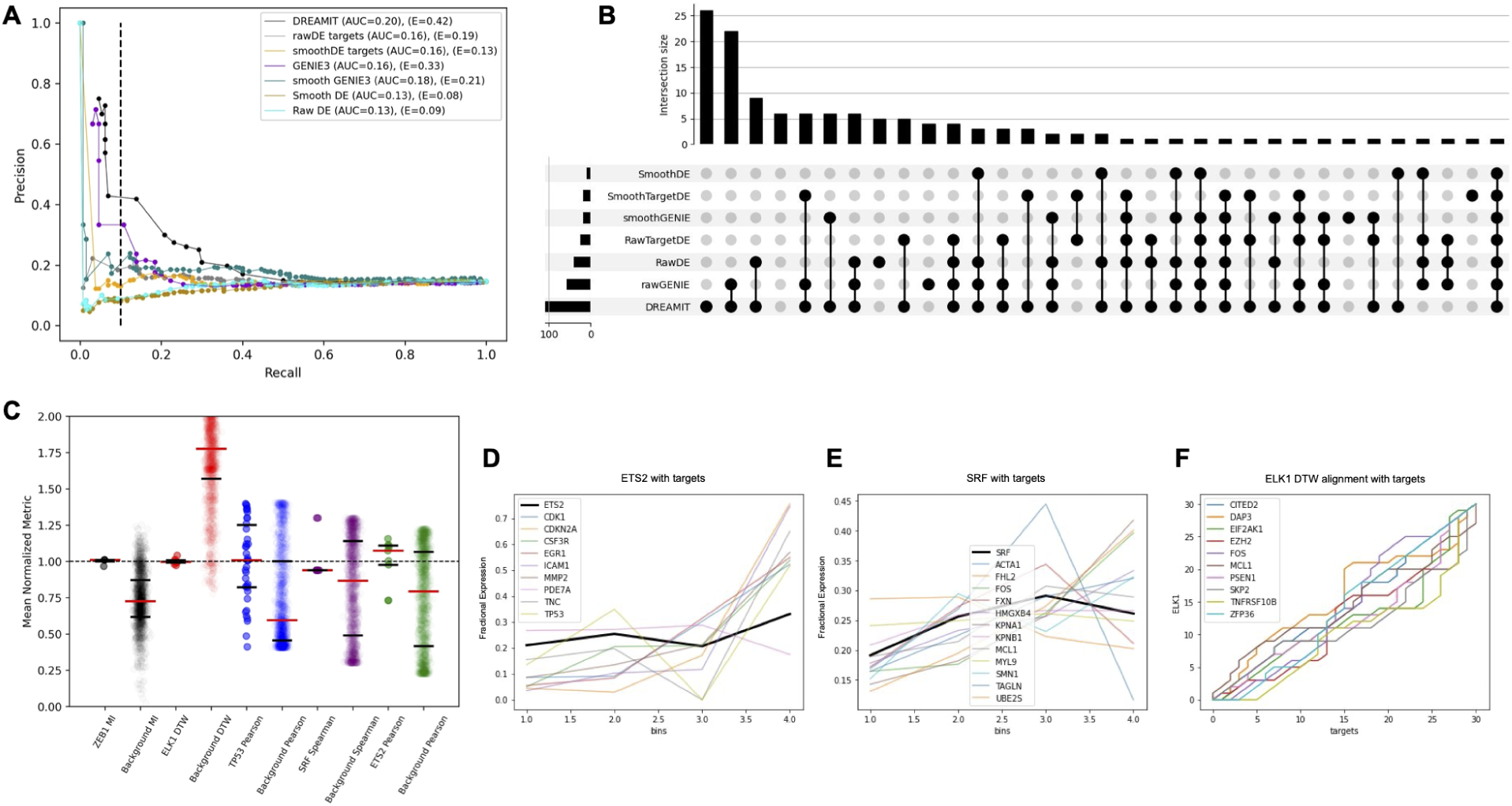
Performance inferring tissue-specific TFs from single cell benchmark datasets. **A.** Precision (y-axis) versus recall (x-axis) measuring each method’s ability to detect TFs from one of the 15 benchmark trajectory branches in which true positive TFs were assumed to be those annotated by TF-Marker DB as previously associated with the tissue assayed by the experiment. **B.** Upset plot illustrating the set intersections between all methods of the TFs inferred for each branch; i.e. set members compared are TF-branch pairs. DREAMIT had the highest number of TF-branch pairs (109) with 26 uniquely predicted. **C.** DREAMIT compares the TF-target relationship distribution to a background for each constituent metric and reports significant TFs through a Kolmogorov-Smirnov test. Examples of five different TFs with different metrics plotted along the x-axis (colors indicate TFs); each TF plotted as a pair with left side showing the observed metric for the targets of the factor and the right side showing the distribution of randomly selected targets. **D**. Illustration of ETS2 factor with its targets found to be significant by Pearson. To visualize genes of different expression scales in one plot, each gene’s expression from one bin was divided by that gene’s expression summed across all bins (fractional expression; y-axis). **E.** Illustration of SRF found to be significantly correlated with its targets using Spearman correlation. Fractional expression (y-axis) was used for visualization purposes. **F.** Dynamic time warping picked up on a significant relationship between ELK1 (y-axis) and its targets (x-axis); the alignment graph illustrates that the targets all maintain a relationship with the factor but there is variability from one target to the next in terms of the exact nature of the relationship.

The DREAMIT component methods (excluded from Figure 3A for clarity) maintained their respective specificity rankings when assessed by precision-recall with the exception of MI (AUC=0.17) and Rolling (AUC=0.18), which both fall behind Pearson (AUC=0.22), Spearman (AUC=0.21), and DTW (AUC=0.19). On the other hand, the early precision of these component methods is ranked quite differently with Rolling (E=0.43) scoring the best (even slightly superior to the DREAMIT ensemble, E=0.42) followed by MI (E=0.38), DTW (E=0.35), Spearman (E=0.28), and Pearson (E=0.28), respectively. Thus, either one of the component methods or the ENSEMBLE as a whole outperformed both DE or GENIE3 in average and early precision, suggesting DREAMIT performs well in detecting the top-ranked TFs as well as overall.

To further compare DREAMIT, DE, and GENIE3, we created an upset plot to view the distinct and common TF-to-branch association pairs found across all of the 15 branches in the benchmark (Figure 3B). The tissue-specific TF-to-branch predictions found at FDR<0.05 in each method are shown. DREAMIT finds the most number of TF-to-branch associations (109; 7.3 TFs per branch) followed by rawGENIE (58; 3.9 TFs per branch), rawDE (40; 2.7 TFs per branch), rawDEtargets (25; 1.7 TFs per branch), smoothGENIE (18; 1.2 TFs per branch), smoothDEtargets (17; 1.1 TFs per branch) and smoothDE (10; 0.7 TFs per branch). DREAMIT produced the largest number of TF predictions per trajectory branch making it the most sensitive method. In addition, DREAMIT shares the most overlap with rawGENIE, 52 associations (89.6%), and with rawDE, 33 associations (82.5%). We also investigated the overlaps of TF-to-branch associations found by the components of DREAMIT (data not shown). DTW found the most associations (73) followed by MI (71), Rolling (66), Pearson (64), and Spearman (39). There was a high degree of overlap amongst all of the methods. Spearman had the least overlapping associations, but 100% of its findings were also reported in one of the other 4 methods. Mutual information and DTW had the most exclusive TF-to-branch associations. Overall 16.5% of the associations found by DREAMIT were found by all component methods, and 66.2% were found by at least two.

Taken together, the vast majority of tissue-specific TF-to-branch associations found by the two competing methods were also found by DREAMIT, and in addition DREAMIT found more than 50 associations missed by these methods. The total number of associations that could have been reported in this analysis was 130. This means that DREAMIT found 83.8% of tissue specific associations, while rawGENIE and rawDE found 44.6% and 30.8%, respectively. DREAMIT had the highest degree of specificity and the highest percentage of tissue-specific markers.

DREAMIT found several cases, missed by other methods, in which the TF-to-target distribution was distinct from the background and found to be significant with a KS test (Figure 3C). First, the marker ETS2 (Figure 3D) was found to be significant according to the DREAMIT Pearson method (Rsq=0.77, FDR<0.001), as well as the marker SRF (Figure 3E) on the same cardiac trajectory branch by the DREAMIT Spearman method (Ssq=0.70, FDR<0.005) [27].

DREAMIT captured these two annotated, biological markers in the trajectory data, while all other methods overlooked this as significant with the exception of rawDEtargets for ETS2 (pval<0.005). Second, the marker TP53 (data not shown) was observed to be significant by DREAMIT Pearson (Rsq=0.77, FDR<1e-7) [28], despite the large number of targets (33) introducing potential noise into the calculation of the Pearson correlation. This demonstrates that DREAMIT is able to find strong TF-to-target relationships in both small and large target sets. This finding was missed by rawDE, rawDEtargets, smoothDE, and smoothGENIE, but was reported to be significant by rawGENIE and smoothDEtargets.

The DREAMIT DTW method also detected a significant association for ELK1 (D=0.24, FDR<0.01) (Figure 3F), an association that was missed by all other methods [27]. Finally, the DREAMIT MI method found significance in the ZEB1 marker (MI=2.35, FDR<0.005), which was missed by rawDE, smoothDE, rawDEtargets, smoothDEtargets, and rawGENIE, but was found by smoothGENIE, suggesting that the smoothing spline implemented provides benefit to other methods of analysis.

To demonstrate the significance and minimal variability of the above results, their distributions were plotted as a swarm plot. The markers ZEB1, TP53, SRF, and ETS2 were all significantly above the background indicating a strong relationship found through either MI content, Pearson correlation, or Spearman correlation. The marker ELK1 was significantly below the background DTW distance, which indicates a stronger DTW alignment for ELK1. Altogether, these findings demonstrate that DREAMIT finds true biology under a variety of data conditions, with a variety of methodological approaches, and that these findings are missed by DE and GENIE methods in many cases.

## Discussion

In this work, we presented DREAMIT, Dynamic Regulation of Expression Across Modules in Inferred Trajectories, a novel framework for testing and identifying dynamic gene regulation in TF to target relationships. The method was developed to aid researchers in identifying the gene-gene regulatory relationships governing the state transitions cells undergo that can be detected from associations in their transcriptomes. Previous methods either ignore gene regulation, such as TRADE-seq or PseudotimeDE [11], or were not designed specifically for single-cell trajectories, such as GENIE3 [6]. As such, DREAMIT provides a complementary perspective on interpreting gene regulatory mechanisms from scRNAseq data.

TF-target sets, whether taken from databases or user-supplied, will likely contain targets that are irrelevant for the analysis of a particular tissue and therefore could obscure the ability to reliably detect an association of a TF to a trajectory branch. For this reason, we introduced the use of a Relational Set Enrichment Analysis (RSEA) to detect the significance of a target set’s association relative to a random background that can tolerate such noise in the target set.

Furthermore, we found that using a target focusing step, much like an enrichment analysis can identify a “leading edge” of contributing pathway genes to a differential expression signature, helps boost the RSEA detection signal.

In a benchmark set of data encompassing six tissues, we found that DREAMIT exceeded the performance of Differential Expression (DE) and GENIE3. DREAMIT had both a higher sensitivity overall and captured more tissue-specific markers.In summary, DREAMIT was shown to have higher sensitivity, finding over 80% of the tested markers, with higher ROC, precision-recall, and early precision compared to DE and GENIE3 based methods.

In the PBMC dataset, DREAMIT also had the highest specificity in terms of ROC (AUC=0.66), precision-recall (AUC=0.57), and early precision (E=1.00). On the erythrocyte branch, DREAMIT found 16 significant TFs, 5 of which were known blood markers and 3 were known stem cell markers. On the monocyte branch, there were 13 significant TFs, 6 were known blood markers and 3 were stem cell markers. Of the significant TFs found by DREAMIT that were not established markers in the TF-marker database [13], 9 of them had established or emerging roles in stem and PBMC development in the literature. For example, the finding association of VDR with monocytes was not documented in the TF marker database but has been demonstrated in recent literature [25,26]. Thus, among DREAMIT’s predictions are potential new examples of tissue-specific regulation by novel factors.

Additionally, DREAMIT was able to find 26 TF-to-branch associations missed by any other competing method (Figure 3B). For example, significant marker findings of ETS2, SRF, TP53, ELK1, and ZEB1 were captured by DREAMIT, but missed by the others. DREAMIT presumably can pick up on these overlooked cases presumably because it considers multiple relationship modalities between TFs and their targets (e.g. Pearson, DTW, MI, etc).

One limitation of DREAMIT is that it considers one TF at a time even though it is known that TFs work together in combination. Even so, several significant associations are found by considering individual TFs in isolation. Extensions of the work are possible that could test combinations of TFs. For example, starting with pairwise TF combination detection, one could test the association between two TFs as well as all pairwise associations between the distinct members of their target sets using the same metrics and statistical tests defined in this work. The drawback of the approach would be the difficulty currently in evaluating its success as there is limited availability of datasets in which TF combinations have been annotated as relevant.

While DREAMIT exceeded competitors in our evaluations, all of the methods had low precision. This is likely due to the limited availability of relevant TFs associated to a particular tissue and to specific branches produced by trajectory inference. For example, using TFs from the TF Marker database allowed us to consider multiple datasets for the evaluation, but it assumed TFs annotated to a tissue were relevant for any/all branches in a dataset that assayed a specific tissue. It is certainly possible that other TFs or other biological differences underlie the variation in the observed transcriptomes of individual cells. There is clearly a need for well annotated datasets that can be used as benchmarks for gene regulatory network inference in single cell analyses [12]. As more datasets with multiple data modalities become available (e.g. ATACseq and RNAseq), it will be possible to develop sets of TFs relevant for branches in a more unbiased and systematic fashion.

DREAMIT code is available on Github https://github.com/nathanmaulding/DREAMIT.git.

## Conclusions

In conclusion, we developed and evaluated DREAMIT, a novel framework for investigating dynamic gene regulation in TF-to-target relationships gleaned from cell trajectories inferred in single-cell RNAseq data. DREAMIT was found to outperform baseline approaches in 15 different trajectory branches in a benchmark dataset and the well-characterized PBMC dataset. DREAMIT detected the association of TFs to tissue-specific trajectories in several instances where the association was missed by all other methods, demonstrating its variety of metrics may help it detect some of the dynamic interdependencies preserved in the pseudotime inference provided by the cell trajectory analysis. In conclusion, DREAMIT offers a complementary approach for shedding light on TF-to-TF networks that govern the temporal regulation assayed by emerging single cell datasets.

## Methods

### Datasets and Dependencies

For each individual cell, DREAMIT uses RNA expression data and assignments to branches and positions along branches (pseudotime) inferred by a cell trajectory inference method (Figure 1A). For this work, slingshot and PAGA were used to infer trajectories and infer pseudotime [29,30]. PAGA was used for the Paul et al. dataset [14] since that study published a set of cell clusters that could be used as input to the PAGA method. Slingshot was used for the benchmark datasets (described next) since no prior clustering assignments were available and because Slingshot has performed well in systematic evaluations [1].

The analysis is dependent on pseudotime assignments estimated by trajectory methods [1]. Errors introduced by a trajectory method may impact the ease with which gene regulatory relationships can be identified. For benchmarking, we collected 7 datasets from EBI representing a diversity of tissues (including brain, heart, embryo, retina, bone marrow, testis) [28], a heart development dataset [27], and a dataset of stem to PBMC lineage [10] for testing DREAMIT. Each of these datasets have undergone preprocessing through the standard scanpy pipeline [7].

Traditional methods compare different states on the branch to each other in terms of differential expression, between a “start” and an “end” state (Figure 1B-C). DREAMIT takes a different approach in which the full set of expression values along pseudotime are used to infer a relationship between a Transcription Factor (TF) and its target. To do this, the method uses a set of predicted linkages between regulators and targets. For this study, we used both the human and mouse regulator-target predictions contained in the TRRUST database [31], which contains interactions mined from over 11,000 PubMed articles. To convert mouse regulogs to human, we used the orthology mapping published by the Mouse Genome Informatics (MGI) consortium [32]. These known regulatory datasets are provided with DREAMIT or the user can choose to include their own regulator-target interactions.

### Trajectory Pre-Processing: Pseudotime Focusing, Expression Quantizing and Spline Smoothing

DREAMIT bases its analysis in first producing a smoothed representation of the data through psuedotime focusing and spline fitting. The use of splines to model scRNAseq data along a trajectory branch follows previous work [10,11]. We selected spline models that incorporated both goodness-of-fit and robustness.

The assignments of cells to locations along a trajectory, commonly referred to as pseudotimes, can reflect a fairly irregular distribution in which the density of cells can vary appreciably from one area to the next. This irregularity in cell number along pseudotime could result in only a few cells, or even a single cell, having a disproportionately large influence in correlation or distance calculations made by DREAMIT (described below). For this reason, DREAMIT applies a *pseudotime focusing* step in which it retains the cells that have been assigned contiguous pseudotime values falling within 1.5 times the interquartile range of all pseudotime values of the branch (Figure 4A-B). The outlier cells that could impact detecting robust trends across pseudotime are eliminated from all subsequent steps.

**Figure 4.**
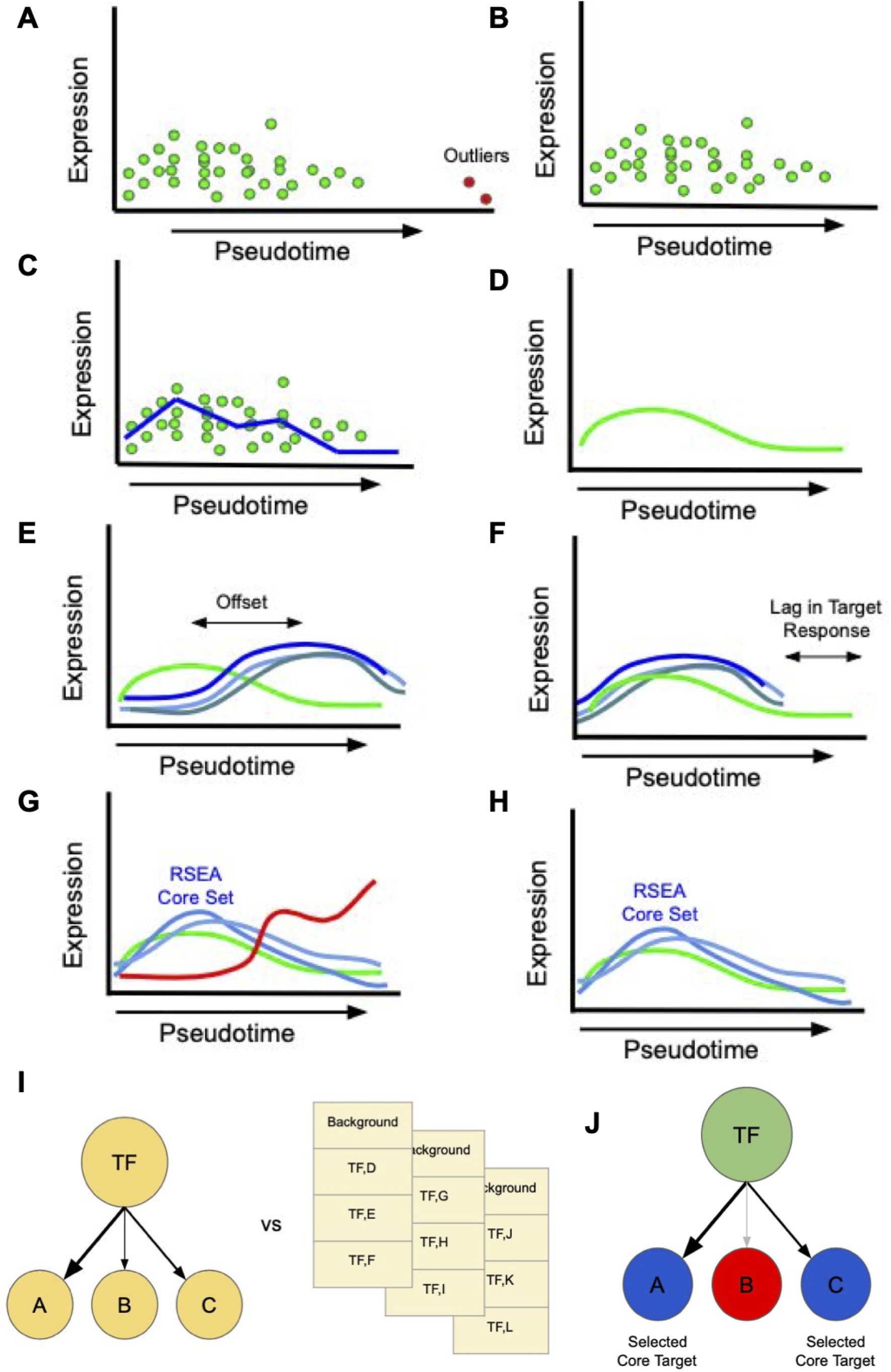
Detecting TF-to-target relations using pseudotime focusing, spline smoothing, target focusing, and significance assessment via random target selection. **A.** Cells with outlier pseudotime assignments (red dots) compared to the other cells (green dots) are shown. B. *Pseudtime focusing* removes outliers from the analysis and retains cells within 1.5 times the interquartile range of all pseudotime values of the branch (see Methods). **C.** Cells are grouped into bins containing at least 10 cells per bin. The average expression of a gene is calculated from all the cells in a bin (blue line) and this bin-averaged expression is used for all subsequent analysis. D. *Spline smoothing* incorporates information from cells in neighboring bins to further smooth out the expression changes in pseudotime (green curve). **E.** DREAMIT quantifies TF-target relationships through pairwise tests of the spline smoothed expression of the TF (green line) and its target genes (blue lines). **F.** Illustration of a “rolling” metric incorporating pseudotime lag. A significant lagged correlation will be detected when several targets share the same delay. G. *Target focusing* employs Relational Set Enrichment Analysis (RSEA, see Methods) to identify a “core” set of targets with high association to the target (blue lines) while excluding the targets with weak or poor association (red curve). H. 75% of the targets with the highest concordance to the factor are retained. I. *Significance is assessed* by comparing TF-to-target metric scores to a random background in which random targets are chosen to be of the same size as the TFs original regulon (yellow lists) with a Kolmogorov-Smirnov test. J. The core set of targets (blue nodes) found by RSEA are used in the statistical analysis.

DREAMIT discretizes the assigned pseudotime values into discrete ordinal levels in order to buffer against further irregularities in the data. Discretization bins are chosen to have an equal number of pseudotime increments, where each bin also has a minimum of 10 cells. Expression levels are then averaged across all cells in the same bin (Figure 4C). Thus, the result of the quantization step produces an average gene expression level for each gene in each ordinal bin. DREAMIT attempts different resolutions of discretization by varying the number of bins from 4 to 100 and chooses a satisfactory number using a hyperparameter search step (described below).

The expression levels of the retained cells could have substantial noise due to technical and biological factors; for example, noise due to the well-documented zero-inflated bias in single-cell RNAseq data [33] (see Figure 5A-B for two examples of raw expression levels). In addition to only averaging the expression levels within a bin, DREAMIT uses a spline smoothing step to incorporate information from cells in neighboring bins (Figure 4D). The discretized pseudotime units are treated as the independent variable and the average expression levels are fit with a zero-inflated negative binomial generalized additive model spline smoothing (NBGAMSS) of the expression values following the recent approaches of TRADE-Seq and PseudotimeDE [10,11]. The number of cells in a bin is used as a "weight" for the fitted data point, placing more emphasis on areas with more support that are assumed to have more reliable estimates of average expression. A smoothing factor determines the number of resulting knots produced by the NBGAMSS (see Figure 5A-B for two examples of smoothed expression). Note that other smoothing operations are possible such as the kernel smoothing used by SINGE [12].

**Figure 5.**
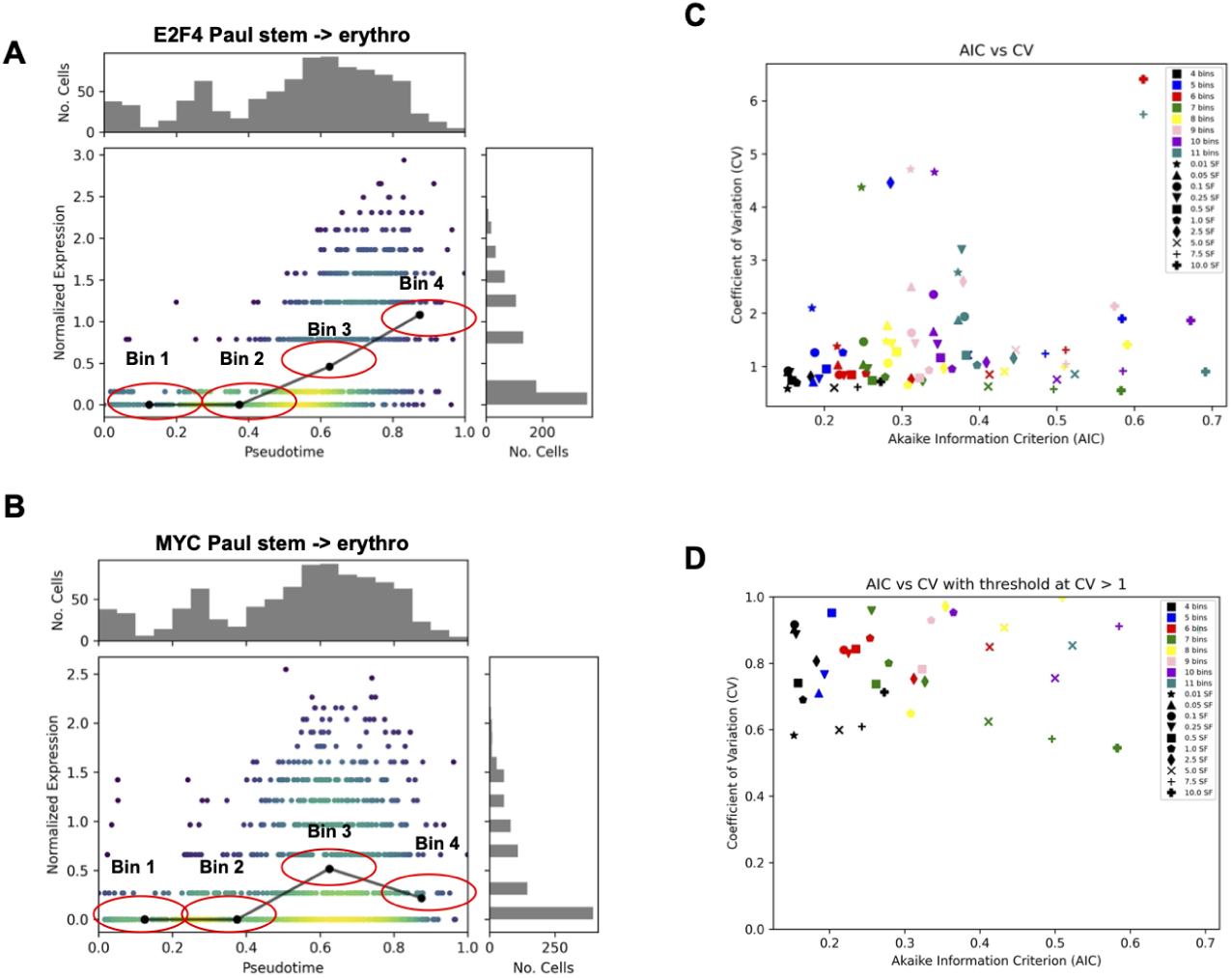
Modeling the gene expression across a trajectory. **A.** Illustration of E2F4 expression (y-axis) across pseudotime (x-axis) along the stem to erythrocyte trajectory branch from the PBMC dataset with normalized (Scanpy) expression (color indicates number of cells); spline-smoothed expression shown for each pseudtime bin (black line); red ellipses illustrate bin width. Histograms plot the distribution of cells at given expression increments (y-axis) and psuedotime increments (x-axis). **B.** Same plot as in A but for the MYC transcription factor. **C.** Each spline parameter choice produces a different fit to the data. Robustness of the fit was assessed by measuring the coefficient of variation (CV) across subsamples of the data (y-axis, see Methods). Goodness of the fit was assessed using Akaike information criterion (AIC) (x-axis). Parameter values that produce spline fits plotted toward the origin (bottom left) are preferred to those further away. **D.** Same as in C but only showing parameterizations that achieve tolerable levels of robustness (CV > 1).

The hyperparameters for a spline are selected by searching for a combination that maximizes the goodness and robustness of the fit, measured by Akaike Information Criterion (AIC) and Coefficient of Variation (CV), respectively. Subsamples are used to compute the Coefficient of Variation (CV) to reflect robustness. We subsample a trajectory branch by choosing without replacement 80% of the cells repeated 30 times. The average Akaike Information Criteria (AIC) is calculated across these subsamples to reflect the accuracy of the fit to the quantized data. Likewise, the CV is calculated across these subsamples. We search for a set of tolerable spline parameters – the number of bins and the smoothing factor – that minimizes both the CV and the AIC. A perfect dataset would have an AIC and CV of 0. We used the distance to the origin of a spline’s associated AIC-CV pair as a measure of goodness that combines both the fit and robustness. To illustrate the spline selection process qualitatively, we plotted splines for ANKRD43 and FRY1 that had a good fit (low AIC and CV; suppl. Fig 1-2) as well as a poor fit (high AIC and CV; Suppl. Fig 3-4). The differences in a poor modeling are evident at high bin numbers, but it should be noted that most spline fits are not extremely off base in terms of modeling, but the best parameters enhance the smoothness. All of the tested spline representations of the PMBC dataset [10] were plotted to assess their AIC (goodness of fit) and CV (robustness of fit)(Figure 5C), with little to no trend being observed with the changing bin and smoothing factor parameters.

Expression data for each of the 100 most variable genes is used for the hyperparameter search. Once an optimal value for these parameters are found, the preprocessing is applied to all genes. In order to ensure that DREAMIT only ever reports on robust representations of the data the maximum threshold for Coefficient of Variation is set to 1 (Figure 5D) in this and all other data analyses. Only trajectory branches that meet this CV criterion will be assessed by DREAMIT.

### Transcription factor to targets

To quantify relationships between a transcription factor (TF) and its targets, DREAMIT performs a series of pairwise tests (e.g. Pearson correlations or dynamic time warping alignments, see next section) between a particular TF and each of its predicted target genes. The targets of a particular TF are taken from the regulatory interactions recorded in the TRRUST database [31].

### DREAMIT metrics to detect a variety of TF-target relationships

The relationship between a TF’s expression levels and that of its target genes could be linear and obvious or it could be subtle and nonlinear, depending on the dataset and the particular regulon in question. For that reason, DREAMIT includes several metrics to pick up on TF-target associations. For each TF with an associated target set, DREAMIT calculates Pearson correlation, Spearman correlation, Dynamic Time Warping Cost, and Mutual Information content [34–37]. We describe the details of the calculation of each of these methods in the following paragraphs.

Pearson correlation provides evidence regarding the strength of the linear relationship between the factor and its targets, while Spearman correlation reflects the strength of the monotonic relationship. Because the factor and targets can have positive or negative correlation, the metric must be squared to create a distribution that is comparable to a random background distribution. Therefore, it is primarily the strength of the relationship that is considered in DREAMIT, but all relationships are included in the report for the user to investigate.

Dynamic Time Warping (DTW) measures how well the expression pattern of a factor can be aligned to the pattern of a target. Target genes may follow a similar pattern as a regulator but have delayed timing that may be detectable in a single cell dataset with enough resolution to reveal cellular processes. DTW finds an optimal match between the patterns that introduces the fewest delays in time [38]. Therefore, a factor with a small DTW distance from its targets implies that the targets are tightly controlled by the transcription factor with few delays in pseudotime.

Mutual Information (MI) describes how much factor expression informs (or could be used to predict) the expression of a target gene. This can be useful for trajectory analysis because it quantifies the information transmission between sender (factor) and receiver (target) [39].

In addition to the standard use of these metrics, a rolling metric calculation is also performed. In the rolling metric calculation, the possibility of a delay in targets responding to factor expression changes is considered by sliding the window in which the relationship is measured. In other words, a rolling window is applied to the target’s expression where each window interval is a bin of smoothed expression (Figure 4E-F). Therefore, the rolling metric reports the bin increment delay at where the peak strength of the delayed relationship occurs. DTW allows for delays in factor target relationships, but is flexible as to the synchronization. In contrast, the rolling metrics report significant factor target relationships with the same time delay, so that all targets respond to the factor expression at a similar pseudotime.

### Relational Set Enrichment Analysis (RSEA): Detecting significant TF-target associations

In addition to a standard reporting of these prior known TF-target associations, DREAMIT uses a two-step Relational Set Enrichment Analysis (RSEA) to highlight the core set of relationships (Figure 4G-J). RSEA begins by performing target focusing to capture the enriched target response followed by a Kolmogorov–Smirnov test (KS test) to assess significance. A transcription factor may activate a subset of its targets in a particular tissue.

DREAMIT attempts to identify the set of utilized targets by identifying a focus set. Target focusing uses the provided set of targets for a factor (e.g. in this study, [31]). DREAMIT performs two steps to identify targets with consistent expression with the factor. First, it checks if the direction of expression matches the activation/inhibition annotation that may be available from the TRRUST database. For example, if TRUST annotates a TF represses a target, DREAMIT checks if the Pearson correlation is negative. Thus, a TF-to-target relation is retained if the sign of the correlation (positive or negative) is consistent with the database annotation, even if the magnitude of the correlation is small. If the database has no annotation about the directionality of regulation (46% of relations in the TRRUST database), then this annotation-expression consistency check is skipped. Second, DREAMIT performs a focusing step to retain a set of targets that mutually have the strongest relations to the TF. Targets with a relationship that falls above the 25th percentile of a DREAMIT metric are retrained in a core “target focus” set. On the flip side, targets with deviant expression are filtered out in an effort to reduce false positives.

To determine the significance of a DREAMIT relationship metric for a TF, the distribution of metric levels between the TF and its targets is compared to a background distribution of the same metric computed between the TF and each of a set of randomly selected target genes. The background set is chosen to be equal to the largest target set considered. After the random targets are selected, the same metrics are calculated for the random set and then compared to the true factor-to-target distribution using the Kolmogorov–Smirnov test (KS test)[40]. A Benjamini-Hochberg correction [41] is then performed on the p-values generated for each factor by the KS test to account for false discoveries. Transcription factors with adjusted p-values < 0.05 are considered significantly non-random for the association in question. In this way, the RSEA results and that of the full set of TF-targets are evaluated, and the core set of relationships is highlighted with target focusing and a KS test.

### Differential Expression (DE)

DREAMIT uses a differential expression (DE) T-test between the start and end of a cell trajectory using the original data values (“Raw DE”) as well as using the spline-inferred data values (“Smooth DE”). The spline-inferred, Smooth DE approach uses a generalized additive model (GAM) as the basis for the DE test, similar to previous methods such as Monocle-2, TRADE-Seq and PseudotimeDE [10,11,42]. For the DE test (either raw or smooth), DREAMIT divides a trajectory branch into two equal segments, the “start” and the “end”. For raw expression, outlier cells are pruned such that those with aberrant pseudotime values are ignored. If a gene shows a statistically significant change between these two conditions then it is considered differentially expressed. Statistics such as fold change, T-statistic, pvalue, and FDR are then reported for each gene on a branch for both the raw and smoothed data.

### Evaluating the specificity of TF to target relationships

To determine the specificity of DREAMIT, ROC, Precision-Recall, and early precision were assessed. True “hits’’ are defined as transcription factors that are markers of the tissue under study. All other transcription factor findings are considered false hits. This assumption does not provide perfect accuracy to true biology as there are likely many factors that have yet to be classified in the TF-Marker database being used [13], and also every factor marker of a tissue need not be active at all times. However, this bronze standard metric does give an enhanced insight into DREAMIT’s ability to highlight tissue specificity. ROC, Precision-recall, and early precision were assessed branch by branch and as an aggregate. These specificity curves were determined for findings by Pearson correlation, Spearman correlation, dynamic time warping, mutual information, rolling calculations, and by the most significant method for each factor. By examining the TPR, FPR, precision, and recall under various p-value thresholds, the specificity of these methods can be assessed comparatively. For early precision, we followed previous works [12] that defined this metric as the precision at recall levels ≤ 0.1.

In a similar manner, competing approaches, differential expression (DE) and GENIE3 [6], were comparatively assessed against these DREAMIT methods. For DE, if the transcription factor (TF) is a listed marker of the tissue under study then it is considered a true hit, while all other TFs are considered false hits. This is the same method used for DREAMIT. Additionally, the DE targets were assessed by comparing the T-statistics of the targets with a background set using the KS test. For the TF in question, it is considered significant if its targets score better than the random background. Both the scanpy processed expression (rawDE) and the spline smoothed expression from DREAMIT (smoothDE) were assessed for both TF DE and DE targets. For GENIE3, a distribution of weights is determined for TFs to known targets from Trrust [31]. This is then statistically compared to a distribution of weights for TFs to randomly selected genes (representing the background) with a KS test. This is done for both scanpy processed expression (rawGENIE3) and the spline smoothed expression from DREAMIT (smoothGENIE3) Specificity is then assessed in the same way as DREAMIT where tissue markers are true hits and all other TFs are false hits.

It should be noted that rawDE has some processing done (IQR pruning of cells with outlier pseudotime assignments). This processing was done, because often dividing a branch strictly on the original pseudotime assignments will result in severe imbalances in the number of cells in the start and end segments. Therefore, some processing was still useful in this analysis above the traditional approach, even before results are considered.

### Evaluating the overlaps between DREAMIT and other methods

In order to compare the findings of DREAMIT, DE, and GENIE3 methods, a set of TF-branch pairs were recorded for each method’s predictions. For example, the pair (MEK,Branch-3) would record that MEK was associated with the third trajectory branch of the dataset. The TF-branch pair sets of each method were compared and visualized using an Upset plot (Fig 3C-D). This was done for both the tissue-specific markers of the respective branches being assessed and for the non-markers for DREAMIT, rawDE, smoothDE, rawDEtargets, smoothDEtargets, rawGENIE, and smoothGENIE. Likewise, the DREAMIT subcomponent methods Pearson, Spearman, DTW, MI, and Rolling were also assessed via Upset plot for both tissue-specific TF marker and non-marker findings.

### Evaluating specificity in reporting TF-markers in a high-fidelity PBMC dataset

To assess the specificity and biology of DREAMIT in a “silver standard”, a PBMC dataset was used [14]. Because this dataset was derived from mice, a set of regulogs [32] was used in this analysis so that downstream specificity and tissue markers can be assessed. Due to a higher proportion of TFs being categorized as blood-specific markers or not, the specificity for DREAMIT can more accurately be determined. ROC, precision-recall, and early precision are determined in the same way as above, using the TF-marker database [13]. Individual factors were then investigated further as to their status in the marker database and in the overall literature.

## Declarations

### Competing Interests Stateme nt

None of the authors have competing interests.

### Data Availability

All data used in this analysis are publicly available

