## Supplementary figures and images for "Associating Transcription Factors to Single-Cell Trajectories with DREAMIT"

### Suppl Figure1

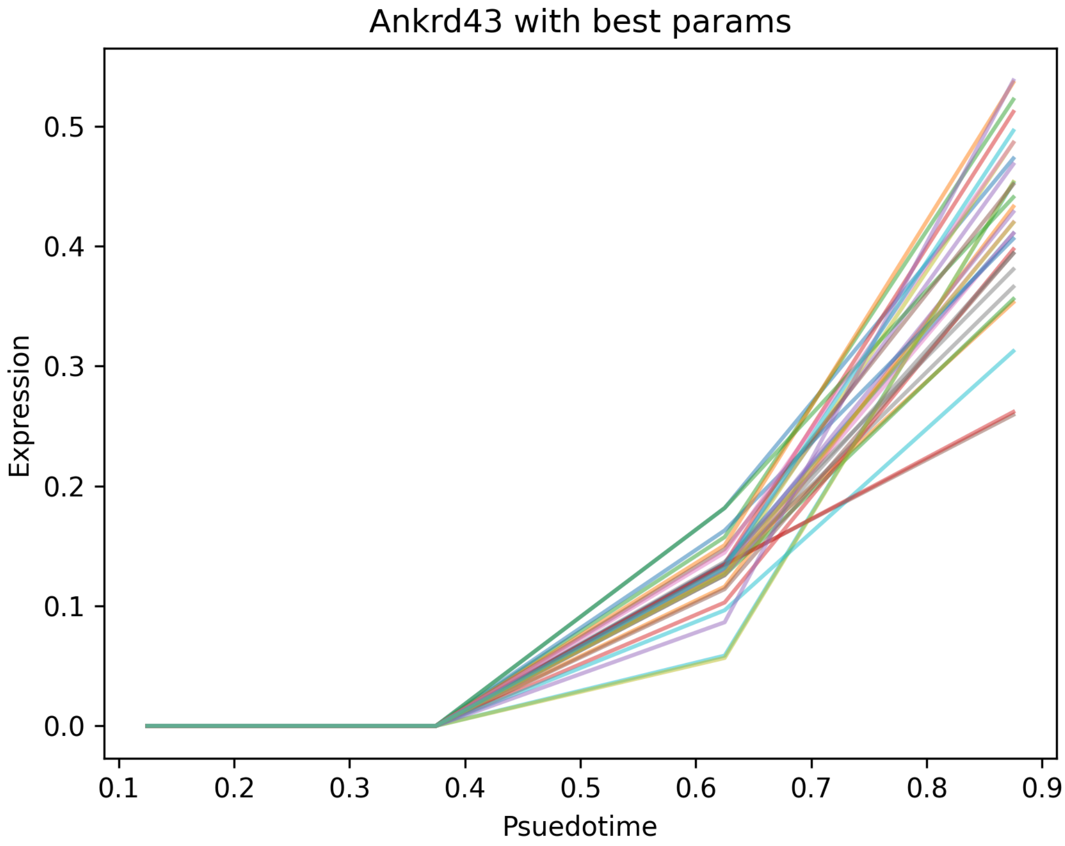

### Suppl Figure2

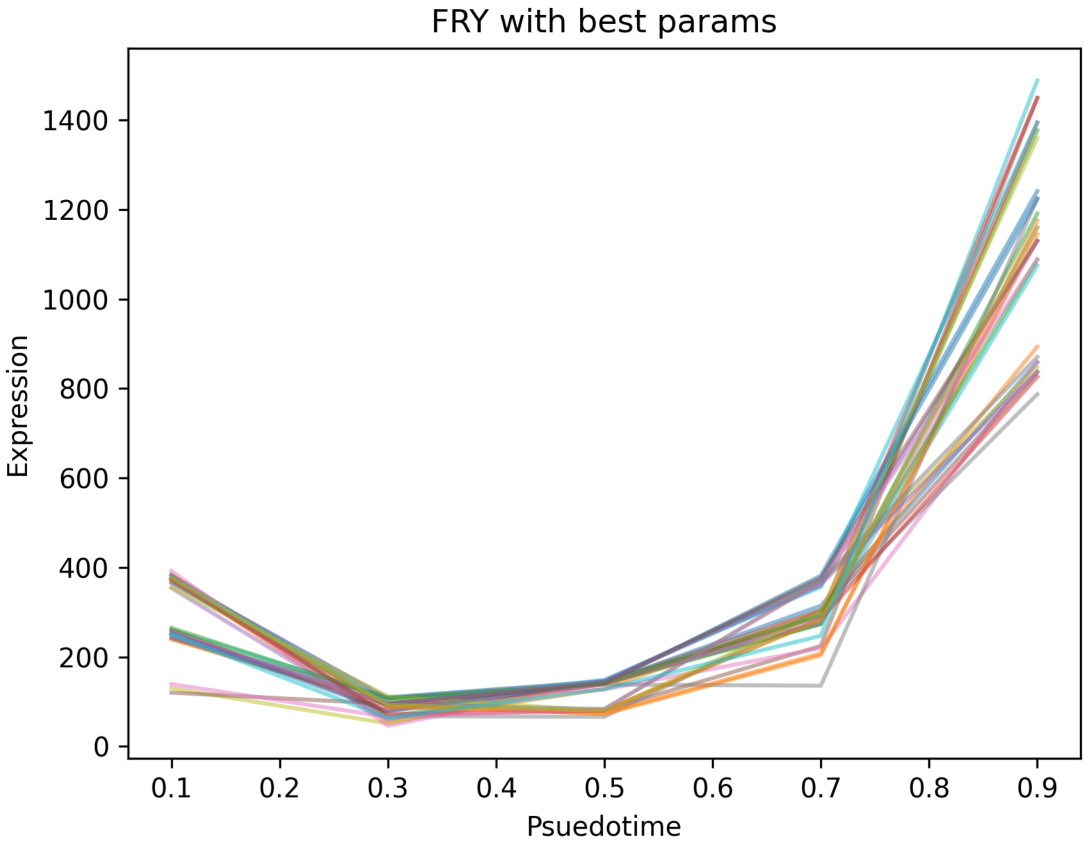

### Suppl Figure3

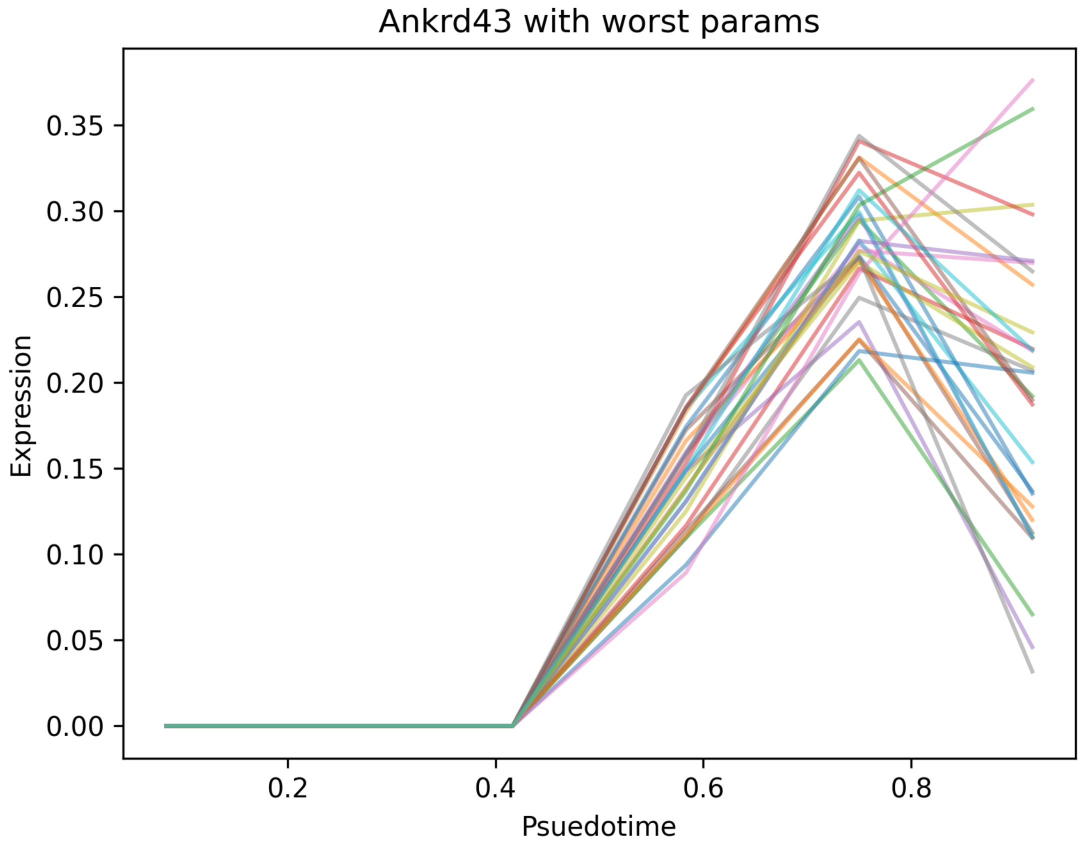

### Suppl Figure4

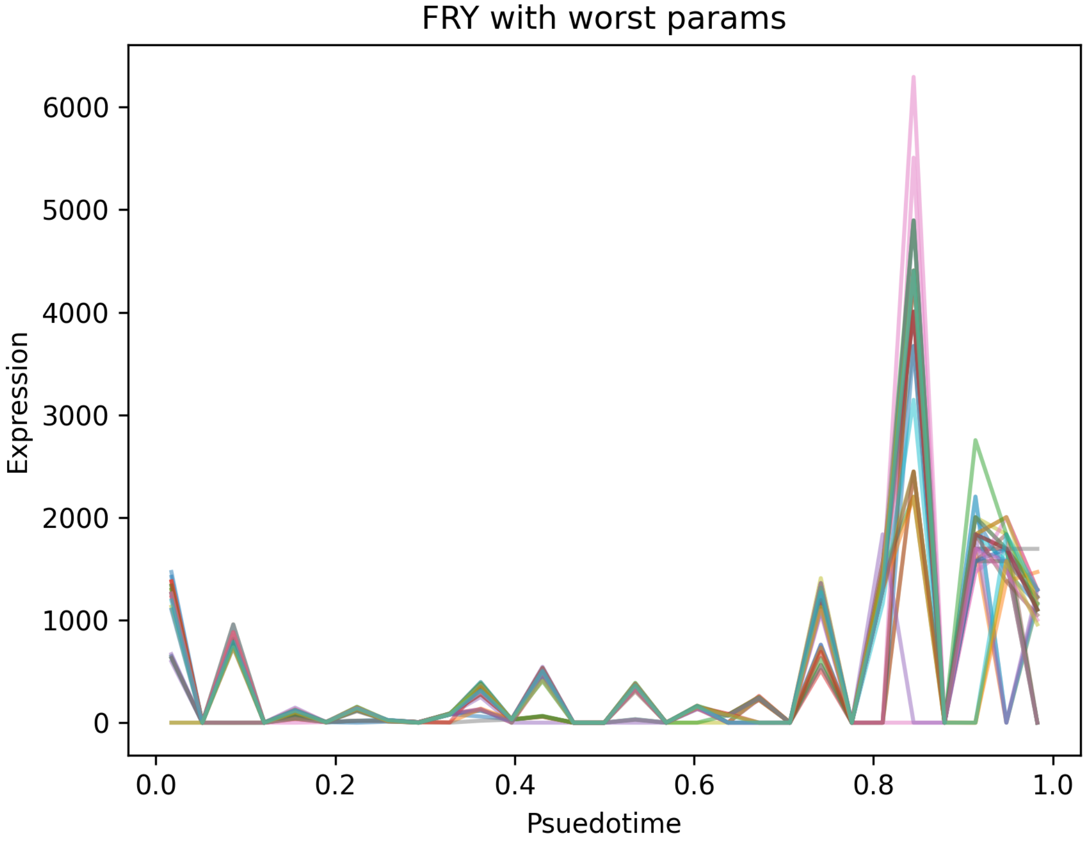
